# Approaches to Species Distribution Modelling in Dinosaurs and Other Fossil Organisms

**DOI:** 10.1101/2023.09.12.557401

**Authors:** Maximilian T. Stockdale

## Abstract

SDM describes a family of methods aiming to predict the geographic distribution of organisms using environmental data such as climate variables. It is a versatile tool in estimating ecological communities and interactions both spatially and through time. However, it has had limited utility in predicting the geographic distribution of fossil organisms. Due to preservation and sampling biases, fossil data does not satisfy the assumptions of species distribution models. Here is presented an analysis proposing a new methodology, which substitutes the conventional pseudoabsence data for fossil occurrences from non-target species. This approach eliminates variation in preservation and sampling, and returns a SDM which performs better than a conventional example. This model concludes that the ceratopsian Triceratops may have been widespread across North America, and beyond the region where its fossils have been recovered. The geographic distribution of Triceratops appears to have been governed by seasonality in temperature and temperature of the coldest month.

## Introduction

Species distribution modelling (SDM) encompasses a range of methodologies which aim to predict the geographic distribution of organisms using environmental data. Such models generalize this task into a classification problem, where a given region on a map is categorized simply as either ‘suitable’ or ‘unsuitable’ according to a selection of environmental variables, usually climate. Species distribution models have seen widespread use in contemporary ecological applications, including conservation biology (Gong et al. 2020, Zhu et al. 2013), rewilding (Sobral-Souza et al. 2017), epidemiology (Escobar and Craft 2016, Escobar 2020), and agriculture (Idohou et al. 2017, Ramírez-Gil et al. 2018, Yang et al. 2020). However, they have seen limited use within palaeontology and for extinct taxa especially.

Previous research has used occurrences of extant organisms, together with phylogenetic bracketing, to extrapolate species distribution models in deep time. Analysis by Watterson et al. (2016) and Harper et al. (2022) trained species distribution models using occurrences of modern reptiles and modelled representations of the current climate. These models were then fitted to palaeoclimate variables derived from a general circulation model, and validated using fossil data. This approach mitigates issues arising from biases in the fossil record. However, it makes a significant assumption that prehistoric examples of a given taxon will occupy the same fundamental niche as their modern counterparts. In addition, this methodology is also limited to taxa which have extant members such as turtles (Watterson et al. 2016) and crocodiles (Harper et al. 2022).

Species distribution models assume that occurrences of a subject organism will be most concentrated in areas where the environment is most amenable to that organism. Fossil data does not fulfil this assumption, because fossilisation takes place under specific environmental conditions. Fossils are expected to be most concentrated where environmental conditions are most favourable to preservation. Fossil occurrences may also be entirely absent from regions that may have been amenable to an extinct organism when it was alive; this is may be a particular issue for areas with mainly low sedimentation rates such as forests.

Absences are an essential point of comparison for SDM . However, demonstrating certain absences is difficult, and finding suitable absence data is an outstanding challenge. Typically such models use so-called “pseudoabsence” data, which draws upon background noise by synthesizing points at random. This approach assumes that a significant statistical signal will emerge from the occurrence data compared to the background. However, this approach requires that the presence data be sampled representatively across the target organism’s geographic range. This is not possible for fossil organisms due to the incompleteness of the rock record. In areas where fossil occurrences are missing, pseudoabsences will still be abundant. This may introduce systematic bias to species distribution models, implying that the environment is more hostile to the subject organism.

A previous analysis (Chiarenza et al. 2019) used an SDM approach to estimate biogeography in non-Avian dinosaurs. This study addressed issues of preservation and sampling biases by reducing the phylogenetic resolution to analyse dinosaurs in general. This reduction in resolution limited the authors to broad-scale macroecological conclusions, and they did not attempt to make genus-or species-level ecological inferences from their results.

An effective methodology for SDM would be an invaluable tool for palaeontologists. In addition to exploring the biogeography of extinct species, such models could be a foundation for understanding other aspects of palaeoecology, population dynamics, food webs, and perhaps even spatial behaviour such as migration. This study proposes an alternative approach to species distribution modelling, optimised specifically for extinct fossil organisms by eliminating variation in preservation and sampling.

## Methods

Two species distribution models were assembled. The initial model used an established conventional methodology to serve as a point of comparison for the second model. A revised model used a novel climate sampling strategy optimized to mitigate some of the impacts of preservation and sampling biases on the fossil record. The dinosaur Triceratops was selected as a test subject. Its fossils are relatively numerous, and originate from sediments in North America which are relatively well sampled (Goodwin et al. 1997). Therefore Triceratops is assumed to be a near-optimal example of a dinosaur fossil record.

The conventional model used fossil occurrence data from the Paleobiology Database (paleobiodb.org). The database was searched for occurrences of Triceratops fossils dated to the Maastrichtian stage of the Late Cretaceous that featured palaeolongitude and palaeolatitude spatial coordinates. This search returned 162 occurrences. This data was shuffled at random and then split into test and training partitions. The training partition included 75% of the original dataset, the testing partition 25%. The conventional model used a pseudoabsence approach to estimate the absence of occurrences. A training sample of 3750 pseudoabsences was generated using the Dismo package in R. A complementary testing sample of 1250 points was also generated. The spatial distribution of these random points was confined to a geographic region corresponding to North America during the Maastrichtian stage of the Late Cretaceous. The palaeogeographic hypothesis was taken from Markwick (2007).

The SDM was trained using palaeoclimate simulations derived from a HadCM3 General Circulation Model. The model parameters are documented in Cox et al. (2001) and Gordon et al. (2000). This model has been used in previous efforts to reconstruct species distribution in deep time (Waterson et al. 2016, Chiarenza et al. 2019, Harper et al. 2022), and therefore offers meaningful comparison. The model output palaeoclimate variables were temperature and precipitation in a monthly resolution. These variables were then processed to derive more relevant bioclimatic variables, include mean annual temperature, mean temperature of each quarter, total precipitation, total precipitation of each quarter, temperature of the warmest and coldest months, precipitation of the wettest and driest months, temperature and precipitation seasonality. These bioclimatic variables were sampled at the x-y coordinates of the presence and pseudoabsence data. The sampled palaeoclimate data was analysed using the Maximum Entropy classifier MaxEnt (Phillips and Dudík 2008). This software has seen widespread use for SDM, and has been shown to outperform other algorithms such as logistic regression (Waterson 2016).

The alternative SDM used the same occurrence data as the conventional model, similarly split into test and training partitions. Pseudoabsence data was not used, instead absences were indicated by the occurrences of fossils from taxa other than Triceratops. The palaeobiology database was searched for Maastrichtian fossils from North America featuring palaeolongitude and palaeolatitude coordinates. The finished dataset included plants, mammals, testudines, crocodylomorphs, lepidosaurs, saurischians, thyreophorans and ornithopods. Other ceratopsians were not included, to limit the biases introduced by any Triceratops specimens which may have been misidentified. The finished dataset included 9200 occurrences, and was split into testing and training partitions. The palaeoclimate data was sampled using the x-y coordinates of all the fossil data, and analysed using a maximum entropy classifier. This SDM was fitted to the palaeoclimate data in the same way as the initial conventional model.

Species distribution models are typically evaluated using an Area Under Curve (AUC) metric. This metric quantifies the performance of a classifier where an AUC of 1 is a perfect classifier, 0.5 is random classification and 0 is a classifier which is entirely incorrect. The AUC metric is estimated using the true positive and false negative rates when the classifier is fitted to a testing partition of the data. This methodology poses this analysis with a dilemma, because the two species distribution models have different associated test partitions, one with pseudoabsences and one with non-ceratopsian fossils. As such the AUC score of the two models would not be easily comparable. To solve this issue, both species distribution models were fitted to both test datasets, returning two AUC estimates per model. These two scores were then averaged to give an estimate of overall performance.

## Results and Discussion

The species distribution models both returned maps of estimated niche suitability for Triceratops (Figure 1). The initial model, estimated using a conventional methodology and utilizing pseudoabsence data, found the potential ecospace for *Triceratops* to be extremely limited and largely confined to the area where its fossils have been recovered (Figure 1A). The revised model, estimated using non-ceratopsian fossil occurrences instead pseudoabsences, found the potential ecospace for *Triceratops* to be very much greater, covering much of the central and southern United States, areas of southeastern Canada and eastern Greenland (Figure 1B). The area estimated to be suitable for Triceratops by the revised model encompasses large areas that are not represented in the Maastrichtian rock record. Superficially this seems more plausible than the initial model, which has returned results that resemble overfitting of the model to fossil data.

**Figure 1:**
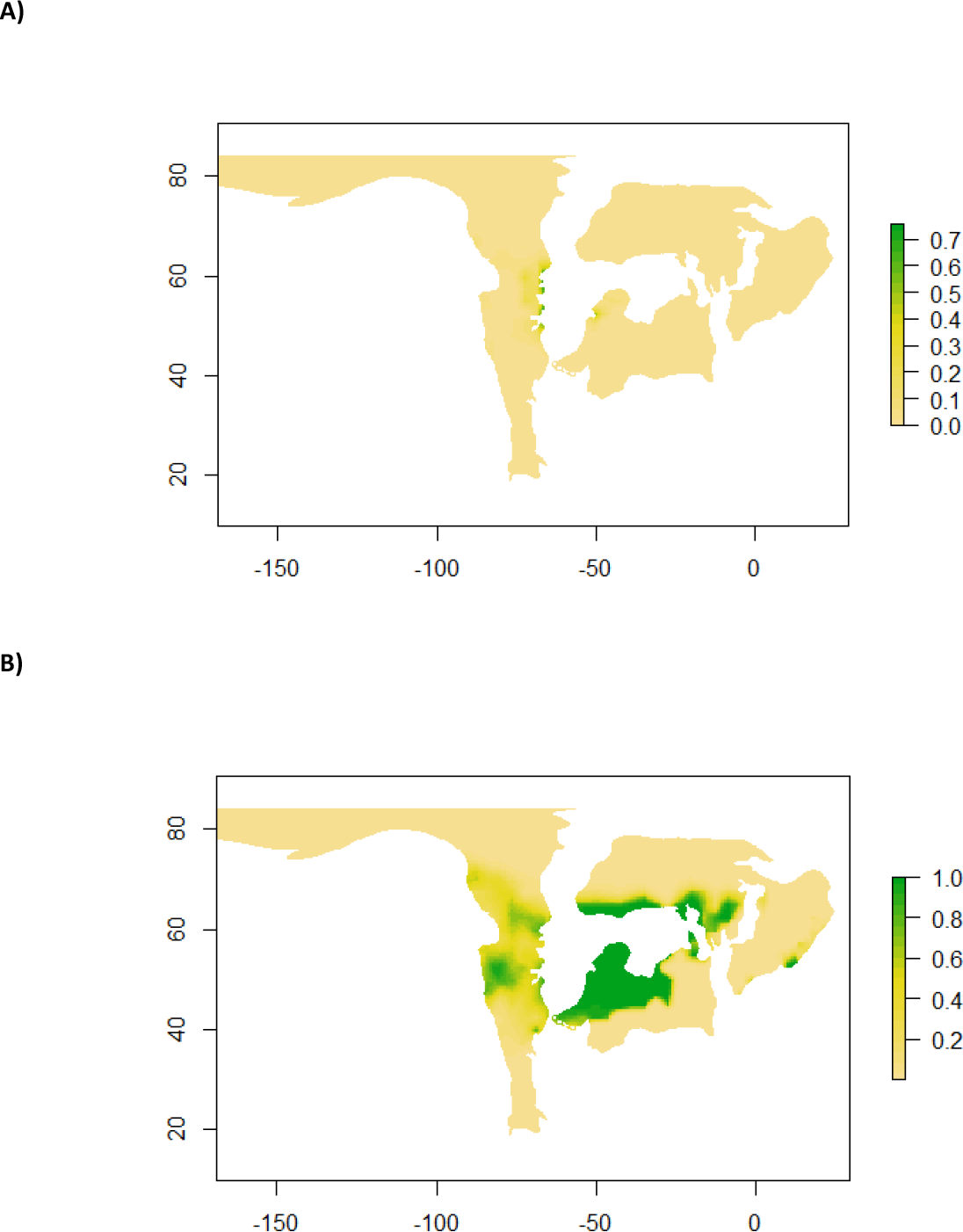
**A)** Map of predicted niche suitability for *Triceratops* according to a species distribution model trained using a combination of fossil occurrences and pseudoabsence data. **B)** A map of predicted niche suitability for *Triceratops* according to a species distribution model trained using a combination of fossil occurrences and occurrences of other species.

The revised model does raise the question of how much of the potential ecospace that was actually inhabited by Triceratops, the “realized niche”. It is possible that all the potential ecospace was occupied, however other factors may have influenced *Triceratops* biogeography. This may have included the distribution of essential food plants, other competing herbivores, environmental variables other than climate, and geographic barriers. The extent of the Western Interior Seaway may have been an obstacle to Triceratops expansion into eastern parts of its potential range. Perhaps the retreat of the Western Interior Seaway in the later Maastrichtian (REFERENCE) opened up potential ecospace to *Triceratops* previously established in Western North America. If the realized niche of Triceratops did not occupy all of the available ecospace, it seems likely that another taxon fulfilled a comparable ecological role in the remaining area. Therefore, species distribution models of other Maastrichtian dinosaurs would be a valuable point of comparison.

The contribution of environmental variables differs between the two models. None of the variables accounted for more than 20% of geographic variation in the initial model (Figure 2). Four of the five highest contributing variables were metrics of precipitation, specifically the first, third and fourth quarter, and the driest month. Most of the temperature variables accounted for less than 1% of geographic variation. The emphasis on precipitation in the initial model is consistent with preservation bias within the fossil data. Areas with high precipitation can be expected to experience more transport and deposition of sediment, through the action of rivers, other surface flows, and flooding. Therefore organisms living in areas of high precipitation may have a greater chance of preservation in the fossil record.

**Figure 2:**
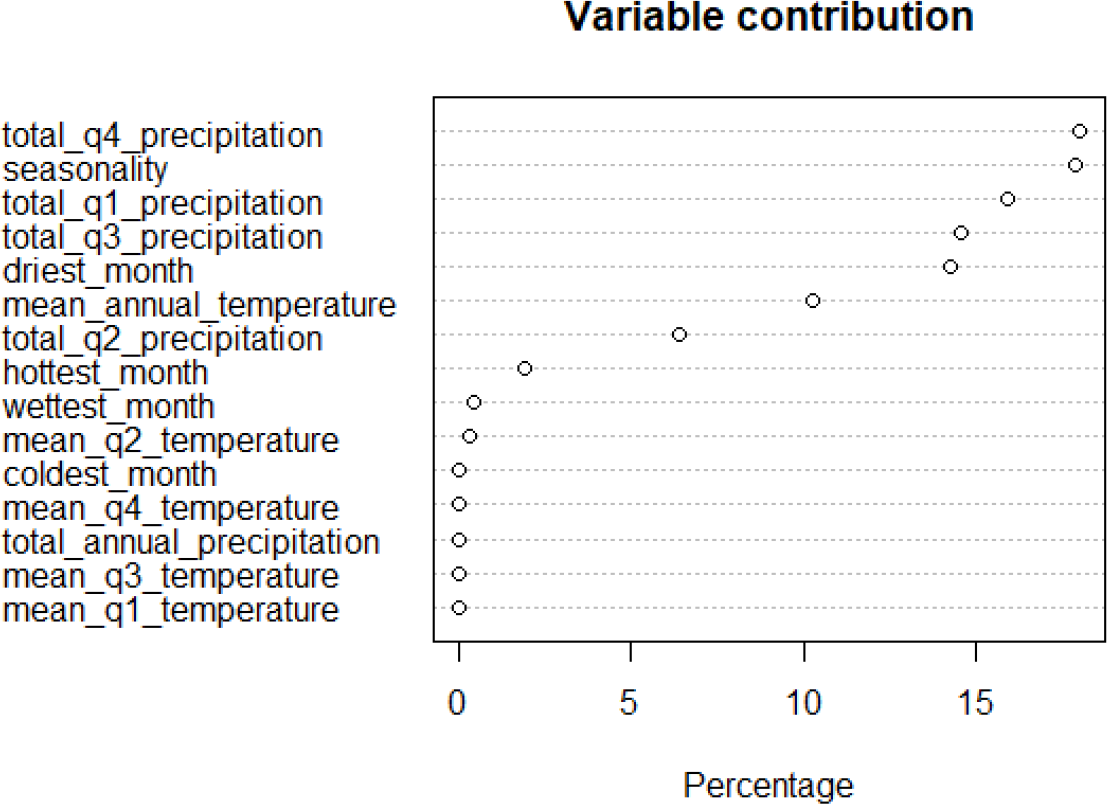
Percentage contributions of environnemental variables to the initial SDM, trained using pseudoabsence data. No variable contributes to more than 20% of geographic variation, with emphasis on precipitation.

The revised model showed a much more significant contribution from temperature variables (Figure 3). The variable with the greatest contribution was temperature in the coldest month, accounting for 50% in variation in geographic distribution. Temperature seasonality also made a large contribution to the model at around 20%. The total precipitation in the first quarter was the precipitation variable with the greatest contribution, accounting for approximately 8% of geographic variation. The revised model shows a much reduced influence from precipitation compared to the initial model. Atmospheric temperature may be expected to vary independently of fossil preservation in a given region. Therefore, the relative importance of temperature over precipitation may be consistent with a decoupling of the model from preservation biases in the fossil training data.

**Figure 3:**
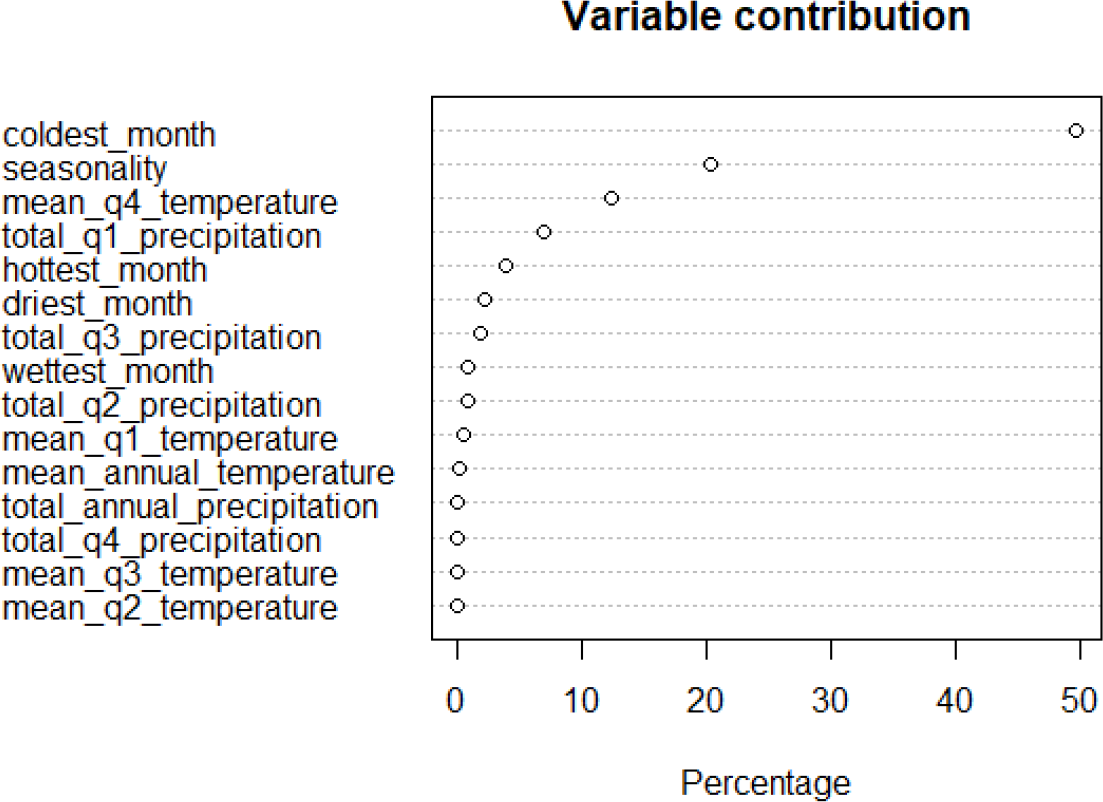
Percentage contributions of environnemental variables to the revised SDM, trained using fossil occurrences to represent absences.

The AUC scores estimated for both models varied depending on what data was used for testing (Table 1). The initial model achieved an AUC score of 0.97 when validated using a testing partition containing pseudoabsence data. Therefore the model was extremely good at distinguishing locations of *Triceratops* fossil occurrences from such pseudoabsences. However, when tested using the fossil testing partition, the AUC score decreases to 0.55 (Table 1). This is close to random classification. Therefore the SDM trained using pseudoabsence data is extremely poor at distinguishing the location of *Triceratops* fossils from those of other organisms. This supports concern that species distribution models of fossil organisms may be susceptible to preservation and sampling bias. It is anticipated that some fossil organisms will exist outside the ecospace available to *Triceratops*. A SDM of *Triceratops* should be capable of classifying these locations in a similar way to pseudoabsences. However, models trained with pseudoabsences seemingly do not have this ability. The revised SDM returned an AUC of 0.76 when it was also tested using fossil data (Table 1).

**Table 1:** AUC scores of the two species distribution models when fitted to different test datasets.

|  |  | Training dataset |  |
| --- | --- | --- | --- |
|  |  | Classic Method<br>(Pseudoabsences) | Novel Method<br>(Fossil occurrences) |
| Testing dataset | Classic Method<br>(Pseudoabsences) | 0.996 | 0.816 |
|  | Novel Method<br>(Fossil occurrences) | 0.55 | 0.762 |
|  | Average | 0.773 | 0.789 |

Therefore this model has a 76% success rate when classifying locations as either suitable for *Triceratops* or non-ceratopsian. This lower AUC score may be the result of a smaller sample size, and the significant geospatial overlap between the Triceratops fossils and the non-ceratopsian fossils.

Perhaps species distribution models trained using fossils to represent absences could be optimized by reducing this relative geographic overlap as much as possible. This could include limiting models to species which are localized in their distribution, or using background fossil data from as wide a geographic area as possible. In addition, it may be possible to use resampling to synthesize bigger datasets which inherit the properties of the fossil data.

The revised model returned an AUC score of 0.82 when tested using pseudoabsences (Table 1). This improvement may be because the pseudoabsences covered the whole of North America, and therefore there was a smaller relative overlap between the presence and absence data in the training dataset. In a sense, perhaps some pseudoabsences are “easier” to classify than most non-ceratopsian fossils, and this might appear to boost the model’s performance. Most importantly, the model trained using fossils performs respectably regardless of whether testing absences are indicated by non-ceratopsian fossils or by random pseudoabsences. This suggests that this revised model is a true SDM of *Triceratops*, and not a distribution model of their fossil occurrences.

On average, the model trained using pseudoabsences returned an AUC of 0.77. The model trained using non-ceratopsian fossils returned an average AUC of 0.79. Therefore, the model trained using non-ceratopsian fossils performs fractionally better.

## Conclusion

A SDM of Triceratops trained using a conventional methodology with pseudoabsence data has limited classifying ability. While it can classify pseudoabsences very well, it has limited ability to distinguish fossil occurrences of Triceratops from that of non-ceratopsians. This suggests systematic bias within the model, since fossil occurrences are likely to occur outside of the ecospace available to Triceratops itself. This model predicted that Triceratops would have an extremely limited geographic range, synonymous with where fossils have been recovered. Precipitation variables contributed substantially to this model, which is consistent with higher rates of transport, deposition and burial associated with conditions for fossilization.

A revised SDM of Triceratops trained using non-ceratopsian fossils instead of pseudoabsences shows respectable performance. This is irrespective of whether this model is tested using pseudoabsence data or non-ceratopsian fossil data. This suggests it is a true SDM of Triceratops, and can predict geographic distribution when fitted to palaeoclimate data. This geographic prediction estimates that Triceratops could have been widespread throughout north America, including large areas that are not represented in the Maastrichtian fossil record. A balanced profile of variables contributed to this revised model, including variables corresponding to temperature, precipitation and seasonality. The model does not place the same emphasis on precipitation as a contributing variable. Instead, the geographic distribution of Triceratops is constrained by temperatures during the coldest month of the year, and by temperature seasonality.

This revised model predicts that the geographic distribution of Triceratops is significantly greater than the geographic distribution of fossils would suggest.

